# Sodium channel slow inactivation normalizes firing in axons with uneven conductance distributions

**DOI:** 10.1101/2022.12.26.521945

**Authors:** Yunliang Zang, Eve Marder, Shimon Marom

**Affiliations:** Volen Center and Biology Department, Brandeis University, Waltham, MA 02454 USA; Technion – Israel Institute of Technology, Haifa 32000, Israel

**Keywords:** Axonal excitability, ion channels, action potential, ectopic spiking, propagation failure, Hodgkin-Huxley model, neuronal resilience

## Abstract

The Na^+^ channels that are important for action potentials show rapid inactivation, a state in which they do not conduct, although the membrane potential remains depolarized^1,2^. Rapid inactivation is a determinant of millisecond scale phenomena, such as spike shape and refractory period. Na^+^ channels also inactivate orders of magnitude more slowly, and therefore have impacts on excitability over much longer time scales than those of a single spike or a single inter-spike interval^3-9^. Here, we focus on the contribution of slow inactivation to the resilience of axonal excitability^10,11^ when ion channels are unevenly distributed across the axonal membrane. We study models in which the voltage-gated Na^+^ and K^+^ channels are unevenly distributed along axons with different variances, capturing the heterogeneity that biological axons display^12^. In the absence of slow inactivation many conductance distributions result in spontaneous tonic activity. Faithful axonal propagation is achieved with the introduction of Na^+^ channel slow inactivation. This “normalization” effect depends on relations between the kinetics of slow inactivation and the firing frequency. Consequently, neurons with characteristically different firing frequencies will need to implement different sets of channel properties to achieve resilience. The results of this study demonstrate the importance of the intrinsic biophysical properties of ion channels in normalizing axonal function.

**Highlights:**

- Variation in ion channel density in axons may compromise axonal spike propagation.
- Slow inactivation of Na^+^ channels modulates their availability.
- Na^+^ channel slow inactivation increases the reliability of spike propagation.
- Normalization by slow inactivation can compensate for uneven channel distributions.

## Results and Discussion

Ion channels have a complex and rich set of slow processes that can capture them in inactivated states, removing them from the easily activated pool for as long as minutes and hours^13-22^. We explore the hypothesis that slow inactivation of axonal Na^+^ channels may be a normalizing factor, contributing to resilience of spike propagation in axons^10,11,23^ with heterogeneous channel densities.

### Slow inactivation

Slow inactivation reflects an activity-dependent transition of an ion channel between two pools of protein states: a pool that is available for activation by membrane voltage within the millisecond time scale of spike generation (depicted *A* in Figure 1A), and a pool that is not available for activation, depicted (*1–A*). The states of a channel in the unavailable pool are coupled by transition rates operating in timescales extending from tens of milliseconds to many minutes. Recent studies demonstrate that at the protein population level, the unavailable (i.e. slowly inactive) pool is a dynamic reservoir that regulates membrane excitability via control of the effective number of ion channels that participate in action potential generation^9,24^.

**Figure 1.**
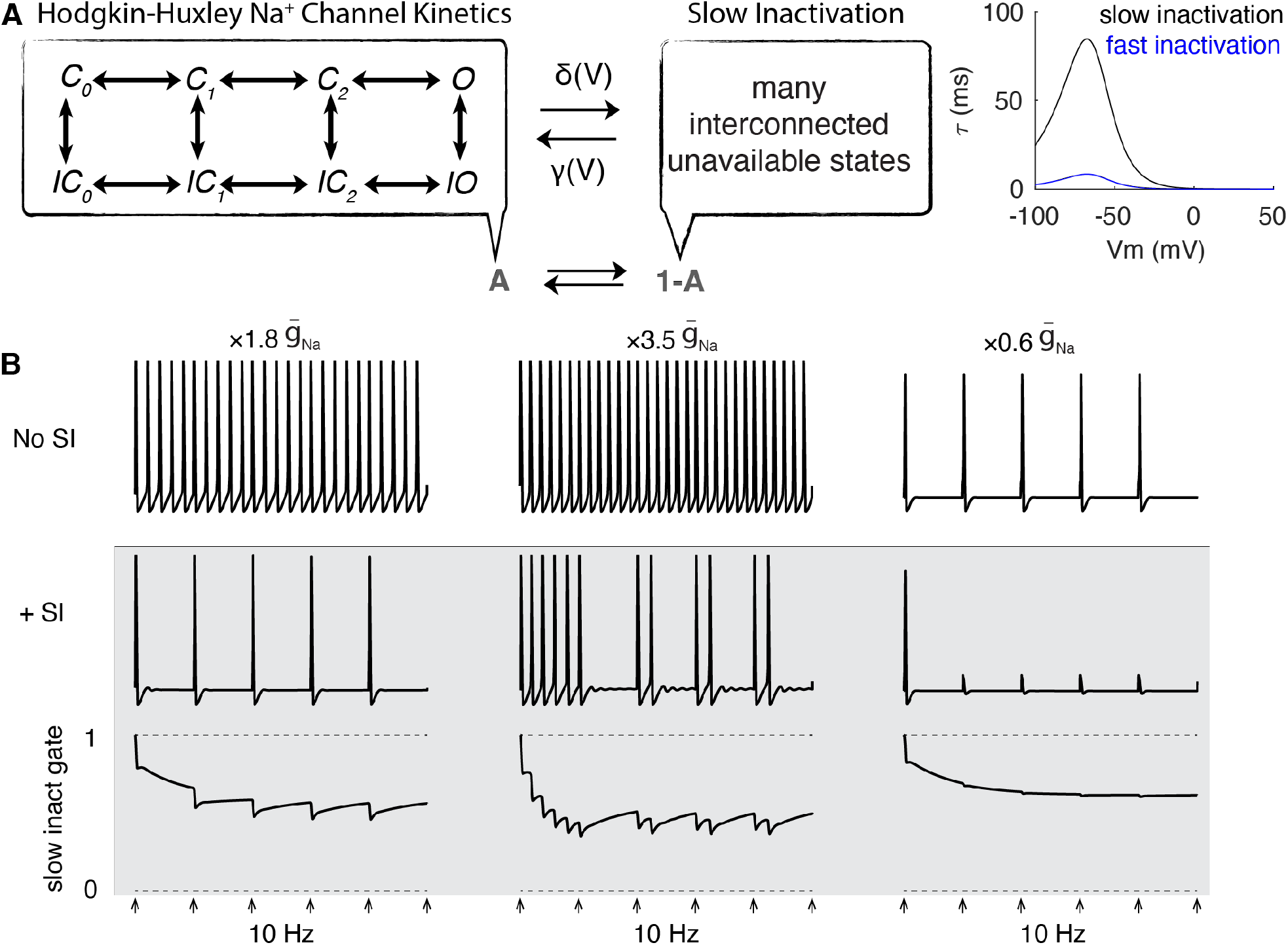
The effect of the Na^+^ channel slow inactivation gate on axonal excitability in a single compartment model. (A) The schematic of Na^+^ channel slow inactivation gate (left) and its time constant (right, black). The blue trace shows the time constant of the h gate in the H-H model. (B) Arrows indicate stimulation times (10 Hz). The top panel has no slow inactivation (no SI). Slow inactivation (+SI) was included in the shaded panels. At 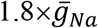 (left) with no SI, the axon fires more rapidly than the stimulation frequency, but with SI, the axon faithfully follows the stimuli (normalization). At 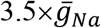 (middle) the axon again fires more rapidly than the stimuli, but the addition of SI produces burst-like interruptions of activity. At 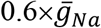 (right) the axon faithfully follows the stimulation frequency in the absence of SI, but with SI it fails to follow repeated stimuli.

### Parameter variability across cells and within a cell

There is a growing appreciation that neurons of the same cell type can be degenerate, that is, produce similar activity with variable sets of conductance densities^23,25-27^. These degenerate solutions arise naturally from the self-organizing homeostatic processes that govern the development and maintenance of the intrinsic electrical properties of neurons^28-30^. Less attention has been paid to understanding the variance of ion channel densities across space in axons. It is likely that ion channel densities are not evenly distributed down the length of axons, and changes in axonal distribution of ion channels undoubtedly occur in disease. The purpose of this study is to understand the tolerance of the action potential propagation to uneven distributions of ion channels, and the role of Na^+^ channel slow inactivation in compensating for variability in channel distributions.

### Na^+^ channel slow inactivation in a single compartment model

The impact of slow inactivation on a single compartment canonical Hodgkin-Huxley (H-H) model^1^ is demonstrated in Figure 1. The slow *A ↔* (*1-A*) kinetics is modelled with forward (*γ*) and backwards (*δ*) rate constants that are scaled versions of the rates governing the standard rapid inactivation in the H-H model. Thus, *δ*(*V*) = 0.1 ∗ *α*_*h*_(*V*) and *γ*(*V*) = 0.1 ∗ *β*_*h*_(*V*) (see Methods). The resulting time constant of the *A ↔* (*1-A*) transition as a function of membrane potential is shown in Figure 1A (right). Figure 1B shows examples of the effects of slow inactivation on spiking in this toy model. As expected, at high maximal sodium conductance the model produces tonic spiking, which may be normalized by the introduction of slow inactivation (Fig. 1B, left). The effect of slow inactivation gate depends on the extent of deviation from the standard H-H maximal sodium conductance (see, for example, middle and right panels of Fig. 1B). Note that while Hodgkin and Huxley’s standard parameter for the sodium conductance is 120 mS/cm^2^, their measured range was 65–260 mS/cm^2^.

### Spike propagation in multicompartmental axonal models with even and uneven distributions of conductances

We built axonal multicompartment models that differed in their Na^+^ and K^+^ channel densities, as illustrated in the cartoon in Figure 2A. In any given compartment, the Na^+^ (blue) and K^+^ (green) conductance densities could be either above or below the values in the H-H model. In these studies, we stimulated the axon at one end at a fixed frequency, and determined the extent to which the entire axon followed the stimulus. Many sets of parameters generated “faithful propagation” (Fig. 2B. left), in that each stimulus generated a propagating action potential. Other parameters resulted in “propagation failure” (Fig. 2B, center). Still other sets of parameters produced “tonic activity” when the axon was active either spontaneously or between stimuli. This occurs when the maximal sodium conductance in one or more compartments is high enough to cause tonic spiking. In the simulation shown in Figure 2B (right), the axon was stimulated at 1Hz at one end, but it was tonically active at ∼ 50 Hz in this case from an ectopic site (high Na^+^ conductance around this location) of action potential generation that triggered spikes that then propagated in both directions.

**Figure 2.**
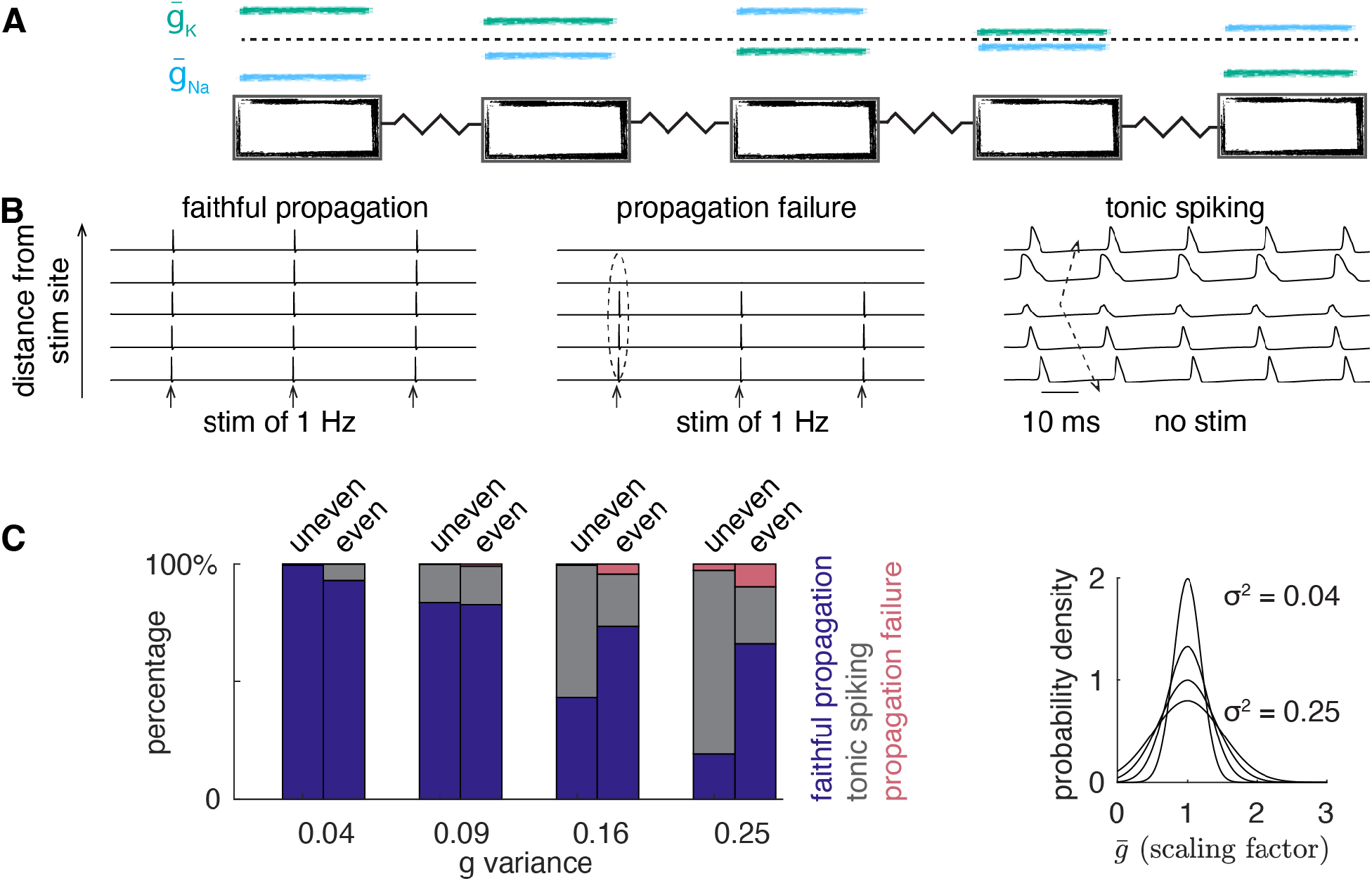
Abnormal spike generation and propagation in axons with varied ratios of channel conductance densities. (A) Schematic of the relative distribution of Na^+^ (blue) and K^+^ (green) conductance densities. The dashed line represents a uniform H-H model. (B) Examples of faithful propagation (left), propagation failure (middle) and tonic spiking (right) in axon models with uneven channel conductance densities. In the left two plots, models were stimulated at 1 Hz, but there is no stimulus during the time range shown in the right plot. (C) The proportion of model behaviors with increased variance of channel conductance densities. For each variance value, the left (right) bar corresponds to axon models with uneven (even) distributed conductance densities. Na^+^ and K^+^ conductance densities were varied by randomly sampling scaling factors from Gaussian distribution functions with different variances (right).

To compare directly the effect of uneven and even channel distributions, we randomly sampled scaling factors for the Na^+^ and K^+^ channel conductance densities (baseline is the H-H model) from Gaussian probability density functions with different variances (0.04, 0.09, 0.16, 0.25). For each variance we simulated a large population of axons (Fig. 2C). The mean of the Gaussian function was 1 (i.e., the H-H value) for all groups (Fig. 2C, right). The histograms in Figure 2C show that at low variances, most axon models showed faithful propagation, but at the lowest variance, the uneven case was more reliable than the even case, presumably because there were axons in which neighboring compartments were able to compensate for compartments with Na^+^ conductances that were insufficient to maintain faithful propagation. As the conductance density variations increased, the number of cases that showed faithful propagation decreased, and the number of cases that showed tonic spiking dramatically increased, especially in axons with uneven distribution of conductances. At higher variances, cases of propagation failure appeared, most notably in the axons with even distributions of conductances.

### Slow Na^+^ channel inactivation in multicompartmental models of axons with even and uneven conductance distributions

Next, we explored whether incorporating a slow inactivation gate can normalize spike generation and propagation by reducing axonal over-excitability. To conveniently manipulate the properties of the slow inactivation gate, we adopted the formulations of Migliore ^31^ (see Methods).

Using this model, the steady state inactivation level, and the time constants of inactivation and recovery from inactivation, can be varied relatively independently (Fig. 3A). In Figure 3B, three types of responses are demonstrated, following introduction of slow inactivation to axon models that produce tonic spiking. Initially the gate was clamped to 1 in all three examples, and they all produced tonic spiking. Once the slow inactivation gate is allowed to change dynamically with spikes, the axonal excitability consistently decreases. If the reduction is in the functional range, tonic spiking will not occur, and axons faithfully spike and propagate only when stimulated (Fig. 3B, top). However, if the reduction is insufficient or excessive, the axons may produce either irregular firing (Fig. 3B, middle) or propagation failure occurs (Fig. 3B, bottom).

**Figure 3.**
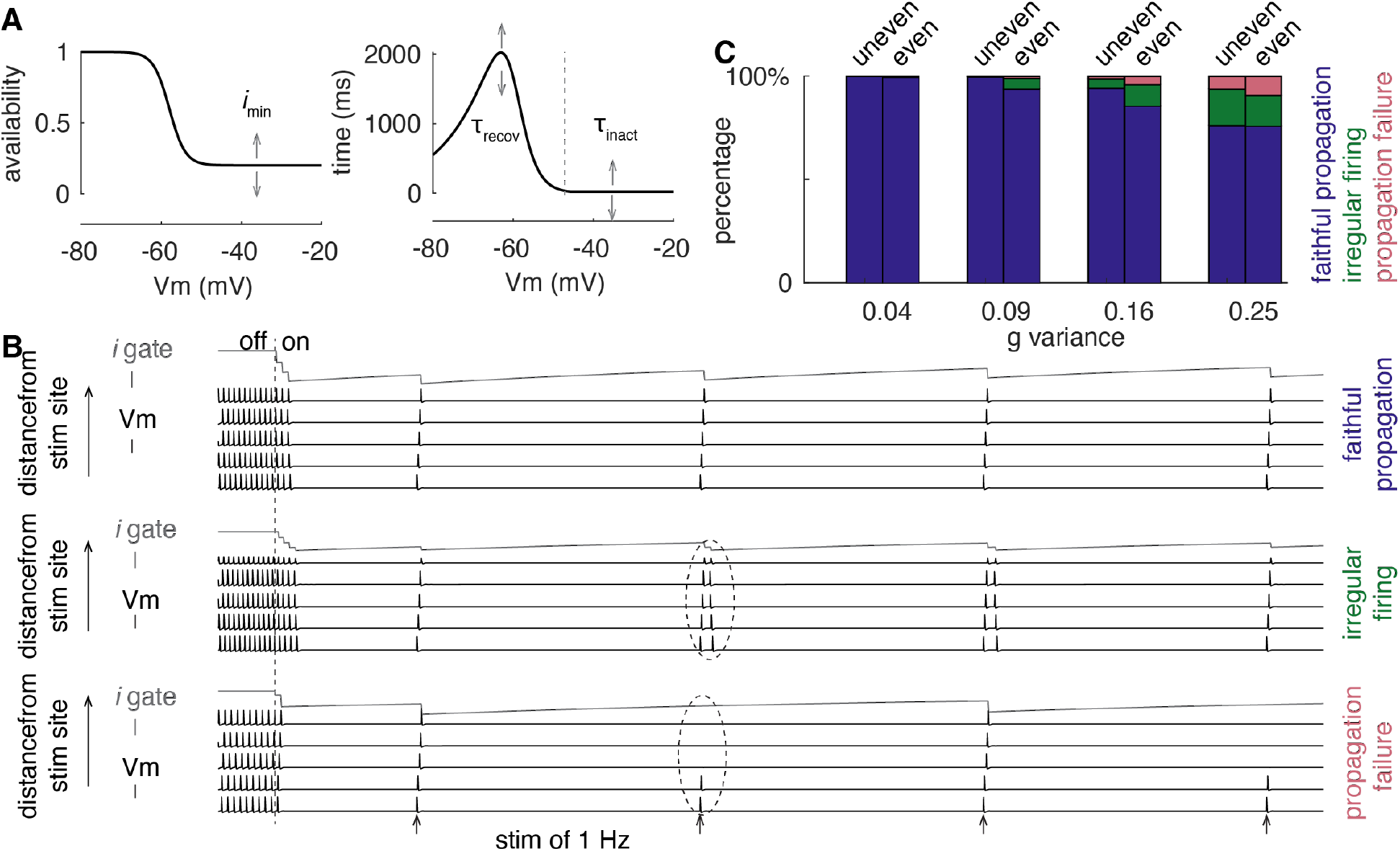
The normalizing effect of the Na^+^ channel slow inactivation gate. (A) The steady state inactivation curve (left) and time constants (right) of the *i* gate. Arrows show parameters whose variations are explored in Figure 4. (B) Typical examples of spike propagation pattern changes before and after switching on the *i* gate. In each plot, black traces represent Vms at different distances from the stimulation site. Gray trace shows the *i* gate at the distally recorded axonal site. Scale bar of *i* gate is 0.6 and scale bar of Vm is 100 mV. (C) The proportion of propagation patterns in axon models with varying degrees of conductance density variances, after including the slow inactivation gate. For each variance value, the left (right) bar corresponds to axon models with uneven (even) distributed conductance densities. Color codes the propagation patterns.

As shown in Figure 3C, incorporating the slow inactivation gate makes most axon models faithfully spike in response to stimuli. By comparison to Figure 2C, incorporating the slow inactivation gate did not substantially improve the propagation reliability of axon models with even conductance density distributions.

### The normalization of spiking by Na^+^ channel slow inactivation is frequency-dependent

We explored how the three parameters (Fig. 3A) of the slow inactivation gate affect its capacity to normalize axonal spike propagation. These parameters represent the steady-state availability of the slow inactivation gate (*i*_min_), the inactivation time constant (*τ*_inact_, time constants at depolarized membrane potentials), and the recovery time constant (*τ*_recov_, time constants at hyperpolarized membrane potentials).

Each model was stimulated at 1 Hz and 10 Hz. The normalizing effect of the slow inactivation gate depends on the stimulation frequency, as seen in the example traces in Figure 4A. This model fired tonically in the absence of slow inactivation. When the slow inactivation was turned on and the axon was stimulated at 1Hz, the model faithfully followed. However, when the same model was stimulated at 10Hz, propagation continued for about the first second, and then it failed. This occurred as a consequence of the slow inactivation that accumulated because of the 10 Hz firing. Interestingly, after about 2 seconds of silence a single spike was propagated, as the inactivation started to wane.

**Figure 4.**
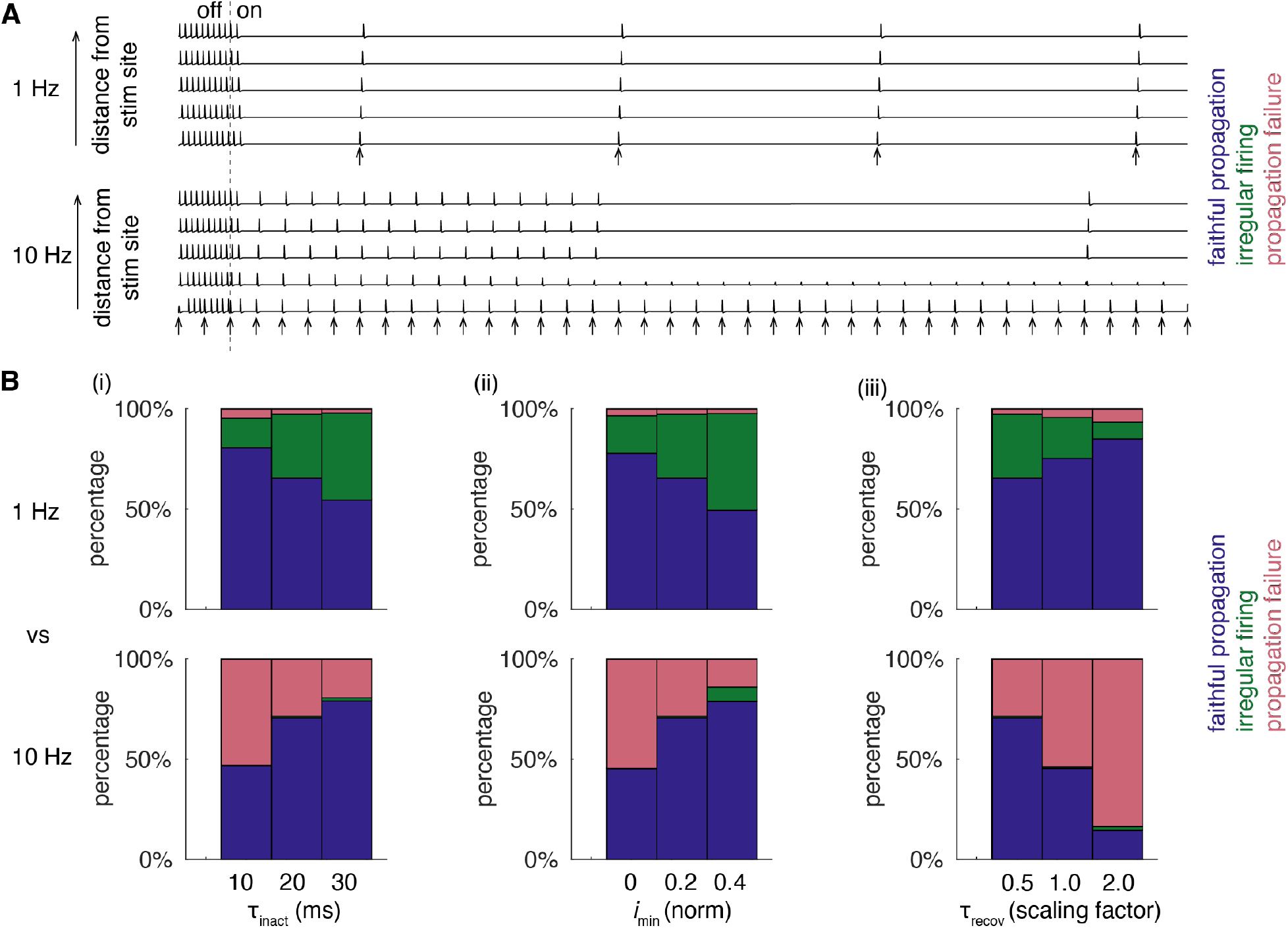
Frequency-dependent normalizing effect of the slow inactivation gate. (A) The effect of the slow inactivation gate on the propagation of spikes at different frequencies. Top to bottom, 1 Hz and 10 Hz, respectively. Traces represent Vms at different sites with distance from the stimulation site. Before the dashed vertical line, the *i* gate was clamped to be 1. (B) The complex relationships among the *i* gate parameters, firing frequency and the normalizing effect. The top and bottom row correspond to stimulation signals at 1 Hz and 10 Hz, respectively. (i) **τ**_inact_ was varied, while *i*_min_ = 0.2 (norm), **τ**_recov_ = 0.5 (scaled); (ii) *i*_min_ (norm) was varied, while **τ**_inact_ = 20 ms, **τ**_recov_ = 0.5 (scaled); (iii) **τ**_recov_ (scaled) was varied, while **τ**_inact_ = 20 ms, *i*_min_ = 0.2 (norm). The axon models have a conductance density variance of 0.25. Color codes propagation patterns.

Figure 4B presents a comparison of the effects of frequency on a population of axons with high variance. The first panel shows that as *τ*_inact_ increases, the rate of faithful spike propagation decreases for stimuli at 1 Hz, but increases for 10 Hz stimulation. The 10 Hz stimulation results in variable amounts of propagation failure, while the 1 Hz stimuli results in a relatively high proportion of irregular firing. Similar results are seen when *i*_min_ is increased. In contrast, when the *τ*_recover_ scaling factor increases, faithful propagation increases at 1 Hz but decreases at 10 Hz.

### Concluding remarks

The frequency-dependent normalizing effects shown in Figure 4 essentially reflect a trade-off between inactivation and recovery of the slow inactivation gate and the chances of triggering irregular firing and propagation failure. This trade-off is activity-dependent, as the time constant of recovery from the unavailable (slow-inactivated) pool is related to the duration of previous activation by a power law^7-9,19,21^.

Different neuronal types show characteristic firing rates, with some that usually fire at very slow rates and others much more quickly^32,33^. One of the implications of this work is that the role of slow inactivation for the normalization of firing rates should match the specifics of the action potential mechanisms and ion channels in each type of neuron. The kinetics of ion channels differ across cell types^34-36^. Thus, cell type definition must also include the specification of the channel properties that can optimally implement normalization by slow inactivation. In conclusion, the effects of variable channel densities across extended morphologies can be complex, and slow channel inactivation is likely to be an important mechanism that is employed by neurons to enhance normalization and thus resilience. Finally, these results demonstrate that improved physiological dynamic range may be a direct result of the properties of proteins, and may not demand regulation mechanisms that involve signal transduction, transcription, and translation.

## Author Contributions

Conceptualization, Y.Z., E. M., and S. M.; Model simulation, Y. Z.; Analysis and Visualization,

Y. Z.; Writing, Y. Z., E. M., and S. M.

## Acknowledgements

E. M. is supported by NIH Grants R35NS097343 and R01MH046742. S. M. is supported by grants from the Israel Science Foundation (ISF 806/19) and the Schaefer Scholars Program at Columbia University’s Vagelos College of Physicians and Surgeons.

## STAR★METHODS

### KEY RESOURCES TABLE

| Software and algorithms |  |  |
| --- | --- | --- |
| NEURON 8.1.0 | Hines and Carnevale, 2006 | https://www.neuron.yale.edu/neuron/ |
| MATLAB R2022a | Math Works | https://www.mathworks.com/products/matlab.html;<br>RRID: SCR_001622 |

### RESOURCE AVAILABILITY

The model simulation code will be public on ModelDB.

## Lead contact

Further information and requests for resources and model codes should be directed to and will be fulfilled by the Lead Contact, Yunliang Zang.

## Method Details

In this work, all the simulations were implemented in NEURON^37^ and visualized in MATLAB. The model formulations were the same as in the original Hodgkin-Huxley model and with a new added *i* gate.

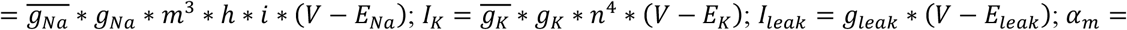;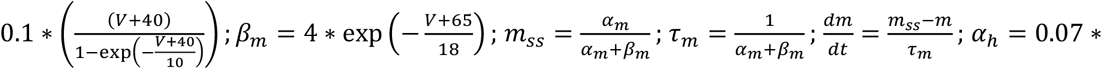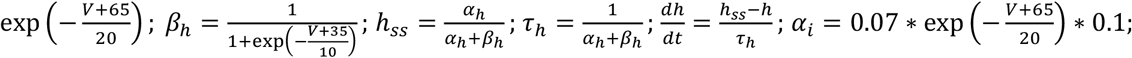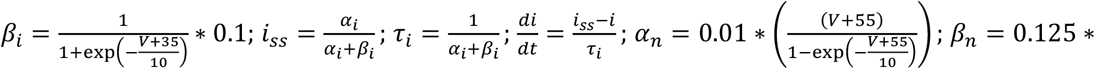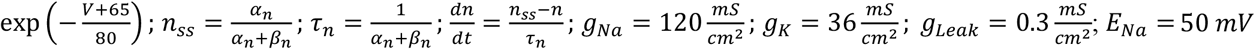 *E*_*K*_ = −77 *mV*; *E*_*leak*_ = −54.4 *mV*. In Figure 1, the slow inactivation gate was simulated by scaling the h-gate kinetics by 0.1. In Figures 2-4, all the formulations are the same except new *i* gate formulations according to the work by Migliore^31^. 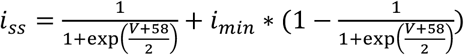; *α*_*i*_ = exp(0.45 ∗ (*V* + 60)) ; *β*_*i*_ = exp(0.09 ∗ (*V* + 60)) ; *τ*_*i*_ = *β*_*i*_/(0.0003 ∗ (1 + *α*_*i*_)); If *τ*_*i*_ ≤ *τ*_*inact*_, *τ*_*i*_ = *τ*_*inact*=_; *τ*_*inact*=_ = 10 − 30 *ms*;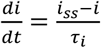; In Figure 4, *i*_*ss*_ was varied by changing *i*_*min*_; the inactivation time constant at high membrane potentials was varied by changing *τ*_*inact*=_; the recovery time constant at low membrane potentials were varied by a scaling factor, because of its voltage dependence implemented in the model. The axon model has a length of 5 mm. To avoid the boundary effect, in the beginning part of the model, we added an axon initial segment with the standard Hodgkin-Huxley model channel densities evenly distributed and the stimuli were exerted at the starting point of the axon initial segment. We only analyzed spike propagation in the first 4.5 mm of the axon to avoid potential boundary effect in the distal end.

